# Drug resistant pancreatic cancer cells exhibit altered biophysical interactions with stromal fibroblasts in imaging studies of 3D co-culture models

**DOI:** 10.1101/2024.07.14.602133

**Authors:** Eric Struth, Maryam Labaf, Vida Karimnia, Yiran Liu, Gwendolyn Cramer, Joanna B. Dahl, Frank J. Slack, Kourosh Zarringhalam, Jonathan P. Celli

**Affiliations:** Department of Physics, University of Massachusetts Boston, Boston, MA 02125; Department of Mathematics, University of Massachusetts Boston, Boston, MA 02125; Center for Personalized Cancer Therapy, University of Massachusetts Boston, Boston, MA 02125; Department of Engineering, University of Massachusetts Boston, Boston, MA 02125; Harvard Medical School Initiative for RNA Medicine, Department of Pathology, Beth Israel Deaconess Medical Center, Boston, MA 02115

**Author notes:** Department of Radiation Oncology, Perelman School of Medicine, University of Pennsylvania, Philadelphia, Pennsylvania (Gwendolyn Cramer).

## Abstract

Interactions between tumor and stromal cells are well known to play a prominent roles in progression of pancreatic ductal adenocarcinoma (PDAC). As knowledge of stromal crosstalk in PDAC has evolved, it has become clear that cancer associated fibroblasts can play both tumor promoting and tumor suppressive roles through a combination of paracrine crosstalk and juxtacrine interactions involving direct physical contact. Another major contributor to dismal survival statistics for PDAC is development of resistance to chemotherapy drugs. Though less is known about how the acquisition of chemoresistance impacts upon tumor-stromal crosstalk. Here, we use 3D co-culture geometries to recapitulate juxtacrine interactions between epithelial and stromal cells. In particular, extracellular matrix (ECM) overlay cultures in which stromal cells (pancreatic stellate cells, or normal human fibroblasts) are placed adjacent to PDAC cells (PANC1), result in direct heterotypic cell adhesions accompanied by dramatic fibroblast contractility which leads to highly condensed macroscopic multicellular aggregates as detected using particle image velocimetry (PIV) analysis to quantify cell velocities over the course of time lapse movie sequences. To investigate how drug resistance impacts these juxtacrine interactions we contrast cultures in which PANC1 are substituted with a drug resistant subline (PANC1-OR) previously established in our lab. We find that heterotypic cell-cell interactions are highly suppressed in drug-resistant cells relative to the parental PANC1 cells. To investigate further we conduct RNA-seq and bioinformatics analysis to identify differential gene expression in PANC1 and PANC1-OR, which shows that negative regulation of cell adhesion molecules, consistent with increased epithelial mesenchymal transition (EMT), is also consistent with loss of hetrotypic cell-cell contact necessary for the contractile behavior observed in drug naïve cultures. Overall these findings elucidate the role of drug-resistance in inhibiting an avenue of stromal crosstalk which is associated with tumor suppression and also help to establish cell culture conditions useful for further mechanistic investigation.

## Introduction

Pancreatic ductal adenocarcinoma (PDAC) is among the most lethal of human malignancies, claiming the lives of more than three quarters of those who are diagnosed within the first year.^1^ Resistance to traditional chemotherapy agents remains a major challenge in clinical management of PDAC. A significant contributor to this is the profound desmoplastic reaction which is characteristic of PDAC. The resultant dense fibrotic stroma is implicated in promoting tumor progression and survival and impeding treatment efficacy.^2–5^ Indeed this has motivated the exploration of stromal depletion treatments, targeting the tumor stroma itself, though with mixed success.^6,7^ Early efforts to therapeutically target cancer associated fibroblasts (CAFs) were promising, but subsequent studies showed that indisciminant depletion of stromal fibroblasts can lead to more aggressive disease progression.^8,9^ Similarly, efforts to improve chemotherapy delivery by targeting Hedgehog pathway signaling yielded promising preclinical results.^10^ However this approach ultimately failed in the clinic and subsequent work indicated that this approach targets stromal elements which otherwise act to restrain PDAC progression.^11^

Stromal involvement in PDAC is complex, with cancer associated fibroblast subpopulations that exhibit both tumor suppressive and tumor-promoting roles. Distinct phenotypic subtypes of CAFs have been identified as inflammatory CAFs (iCAFs) and myofibroblastic CAFs (MyCAFs).^12–14^ In the tumor microenvironment, iCAFs, which are not typically in direct physical contact with cancer cells, activate pathways that promote tumor proliferation and survival via paracrine signaling. MyCAFs interact with epithelial cells through direct cell-cell adhesions, which can play tumor suppressive roles by physically restraining invasive progression. This juxtacrine signaling is mediated by cell-cell adhesions and the genes that code for adhesion molecules, which are also associated with regulation of PDAC progression.^15–17^

In PDAC, as with other carcinomas, drug resistance is one of the most significant barriers to achieving successful treatment outcomes.^18^ Through both intrinsic and acquired resistance, tumors often are, or become, non-responsive to chemotherapy agents, leading to recurrence and relapse even when initial treatment response is positive. Non-responsiveness to therapeutics is compounded by association between drug resistance and epithelial mesenchymal transition (EMT), which acts as a regulator of the tumor-initiating cancer stem cell (CSC) phenotype.^19^ EMT, which is directly linked with drug resistance in PDAC,^20^ also imparts increased metastatic potential.^21^ As such, the tumor cell popualtions which are non-responsive to treatment are also associated with lethal metastatic progression. Less is known however about how the acquisition of chemoresistance alters heterotypic interactions between tumor and stromal cells in the PDAC microenvironment, especially in view of relatively recent knowledge of CAF subpopulations. In the present study we examine the role of drug-resistance using a chemoresistant sub-line previously established in our lab by exposure of PANC1 cells to oxaliplatin chemotherapy over multiple passages. The drug-resistant cell-line PANC1-OR, which has been stable over multiple passages and cryopreservation, exhibits a clinically-relevant oxaliplatin and gemcitabine-resistant phenotype along with increased invasive behavior, and phenotypic traits consistent with increased EMT.^2223^

In this study we use in vitro 3D co-culture models to examine A) the role of co-culture geometry in recreating interactions between adjacent and distant cells, and B) how these interactions become altered when PDAC cells acquire chemoresistance. In both cases we leverage time lapse microscopy and particle image velocimetry analysis (PIV) of co-cultures as a means to observe growth behavior and qualitative biophysical interactions over time. To examine how phenotypic changes associated with drug resistance impact upon tumor-stroma interactions, we correlate imaging data with mRNA sequencing to examine differential gene expression between drug naïve and drug resistant PDAC cells.

In this study we initially contrast two different co-culture geometries as shown in Figure 1. In an embedded-fibroblast-with-overlaid-cancer-cells co-culture (EOC) model, the fibroblastic cells (either Pancreatic Stellate Cells, PSC, or MRC5, normal human fibroblasts) are suspended within a layer of ECM and the epithelial PDAC cells are overlaid above the ECM, allowing biochemical crosstalk without physical contact. We contrast this with an adjacent-overlay-co-culture (AOC) model, where we incubate 3D PANC1 cells for several days to form multicellular nodules, then introduce fibroblastic cells in the same plane on the surface of the ECM bed. This AOC model, which allows for direct physical contact and juxtacrine interactions, is the main focus of the present study. Time-lapse imaging and particle-image velocimetry analysis of cell co-migration and fibroblast contractility in AOCs, measured here as the speed of nodule aggregation, allow for quantitative interrogation of biophysical interactions between tumor and stromal cells for each experimental condition.

## Materials and Methods

### 2.1 Cell Culture

PANC1 and MRC5 cell lines were acquired from ATCC and maintained in T75 cell culture flasks according to ATCC guidelines. DMEM and MEM (HyClone) were supplemented with 10% FBS (HyClone), 100IU/mL penicillin, 1% streptomycin (HyClone), and 0.5 mg/mL amphotericin B (Corning). PSC cells were obtained from ScienCell. SteCM (Stellate cell media, ScienCell) was supplemented with 2% FBS, 1% SteCGS (Stellate cell growth supplement, ScienCell), and 1% penicillin. PSC and MRC5 cells were used for experiments between the 3^rd^ and 10^th^ passages. PANC1-OR is a stable oxaliplatin resistant sub-line of PANC1 developed and characterized previously.^17^ PANC1-OR cells were maintained in T75 flasks according to ATCC guidelines for the PANC1 parent line.

### 2.2 Analysis of mRNA-seq data

RNA was extracted from triplicate cell cultures of PANC1 and PANC1OR cells using the TRIzol reagent (Life Technologies Corporation, Carlsbad, CA). RNAseq was performed by the Center for Personalized Therapy Genomics Core at University of Massachusetts Boston. The quality of the raw fastq files were assessed using FastQC (v.0.11.5).^25^ Adaptor sequences, “AGATCGGAAGAGCACACGTCTGAACTCCAGTCA”, and “AGATCGGAAGAGCGTCGTGTAGGGAAAGAGTGT” were trimmed from the 3’ end of the reads using Cutadapt commandline tool form the Trime Galore package (v.0.4.2). The trimmed reads were mapped against the human reference genome (Ensemble, GRCh38) using STAR/2.5 using default parameters.^26^ The average alignment rate was 96%. The sorted BAM files generated by STAR were used to estimate the transcript abundance per sample using featureCount from the Subread package (v.1.6.2).^27^ Gene expression analysis was performed using the edgR Bioconductor R package (v.3.24.3).^28^ The edgeR TMM method (trimmed mean of M values) was applied to the filtered genes utilizing the DGElist(), calcNormFactors(), estimateGLMCommonDisp(), estimateGLMTrendedDisp(), estimateGLMTagwiseDisp() functions. The glmFit and glmLRT functions from edgeR were used to fit a negative binomial generalized log-linear model to the read counts. The expression of the genes was ranked by logFoldChange (logFC) and false discovery rate (FDR). Differentially expressed genes (DEGs) were determined using abs(logFC) > 2 and FDR < 0.01 cutoffs, resulting in 1342 protein coding DEGs.

### 2.3 Enrichment analysis

Gene ontology (GO) analysis was performed on differentially expressed genes using the g:Profiler with g:SCS (https://biit.cs.ut.ee/gprofiler).^29^ GO terms and KEGG and Reactom pathways with significant overlap with up and down regulated genes were determined (FDR < 0.05) and visualized with Heatmaps using the ComplexHeatmap R package (v.1.20.0).^30^ To run the gene set enrichment analysis (GSEA), we used the fgsea Biocondoctor R package (v.1.9.5).^31^

### 2.4 Spheroid Preparation and co-culturing

Prior to plating cells, GFR Matrigel^TM^ was thawed overnight at 4°C and kept on ice until use. For the adjacent overlay co-cultures, 225ul of growth factor reduced (GFR) Matrigel^TM^ was added to the center of each well of a pre-chilled black-walled 24-well plate (Ibidi USA inc.). After the addition of Matrigel^TM^ the plate was agitated to ensure an even coat on the bottom of the well and then the plate was incubated at 37° C for 20 minutes allowing the Matrigel^TM^ to solidify. PANC-1 cells or PANC-1-OR cells were collected, and cell density was determined using an automated cell counter. Preparations of each line were made at a concentration of 7500 cells/mL of media. 1mL of the PANC1 cell preparation was plated in each of six Matrigel^TM^ coated wells of the prepared 24 well plate. This was repeated with the PANC-1-OR subline. The cells were allowed to incubate at 37°C and 5% CO2 for 7 days. After 7 days of spheroid growth, MRC5 or PSC cell lines were collected and counted. The cells were pelletized and resuspended in DMEM at a concentration of 1 x 10^5^ cells/mL of media. Media was removed from each well containing tumor spheroids. A 1 mL volume of the prepared MRC5-media solution was added to each of three wells of PANC1 spheroids and three wells of PANC1-OR spheroids. One mL of the prepared PSC-media solution was added to each of three wells of PANC1 spheroids.

For the EOC model, MRC5 or PSC cells collected and suspended in Matrigel^TM^ and 225 ul of the fibroblast-Matrigel^TM^ suspension were added to the center of each well. After the addition of Matrigel^TM^ the plate was agitated to ensure an even coat on the bottom of the well and then placed in the incubator until the Matrigel^TM^ set. PANC1 cells were collected, and cell density was determined using an automated cell counter. A preparation of PANC1 cells in media was made at a concentration of 7500 cells/mL of media. A 1 mL volume of the PANC1 cell-media preparation was plated in three MRC5-Matrigel^TM^ suspension coated wells and three PSC-Matrigel^TM^ suspension coated wells. The cells were allowed to incubate at 37°C and 5% CO^2^ for 7 days.

### 2.5 Imaging and image Analysis

Time-lapse 10x phase contrast images and 1.25x bright field images were taken using an inverted time lapse microscope(EVOS, Thermo Fisher Scientific) at a rate of 10 minutes per frame for 4 days. Image data was collected from three independent cell platings containing AOC, EOC and homotypic culture conditions and processed with custom Matlab image processing routines to obtain spheroid size distribution histograms from sets of segmented images. Velocimetric data was obtained using PIVlab v1.43, an open source PIV toolbox for Matlab.^32,33^ For 10x image data pre-processing was done in PIVlab. CLAHE, high pass, intensity capping, and denoise filters were all enabled in order to optimize in-app segmentation for analysis. For 1.25x bright field image data, images werefirst segmented, and preprocessing was disabled in PIVlab. In both cases an 800x800 pixel region of interest was chosen in the middle of the well to eliminate anomalies at the edge of wells. Multi-pass PIV analysis of each experimental condition was completed in triplicate utilizing the fast Fourier transform (FFT) window deformation algorithm generating velocimetric data. Here pairs of images are analyzed in multiple passes. Interrogation windows within the region of interest were cross-correlated in the frequency domain to generate a the velocity vector map of object speed across the region of interest. Object speed for each of the three iterations of each experimental condition was then averaged across each time-matched frame providing a basis for comparing the relative changes in average speed under each experimental condition. Curve Fits were performed using OriginPro software. For statistical analysis of variation in velocimetry profiles between conditions, the mean velocity and standard error across replicates was computed for each timepoint and used to calculate chi-squared values, and ultimately overall p-value for each comparison between conditions based on chi-squared and the number of degrees of freedom.

## Results and Discussion

### 3.1 Imaging-based analysis of PDAC-fibroblast co-culture growth, aggregation and contractile behavior

**Figure 1:**
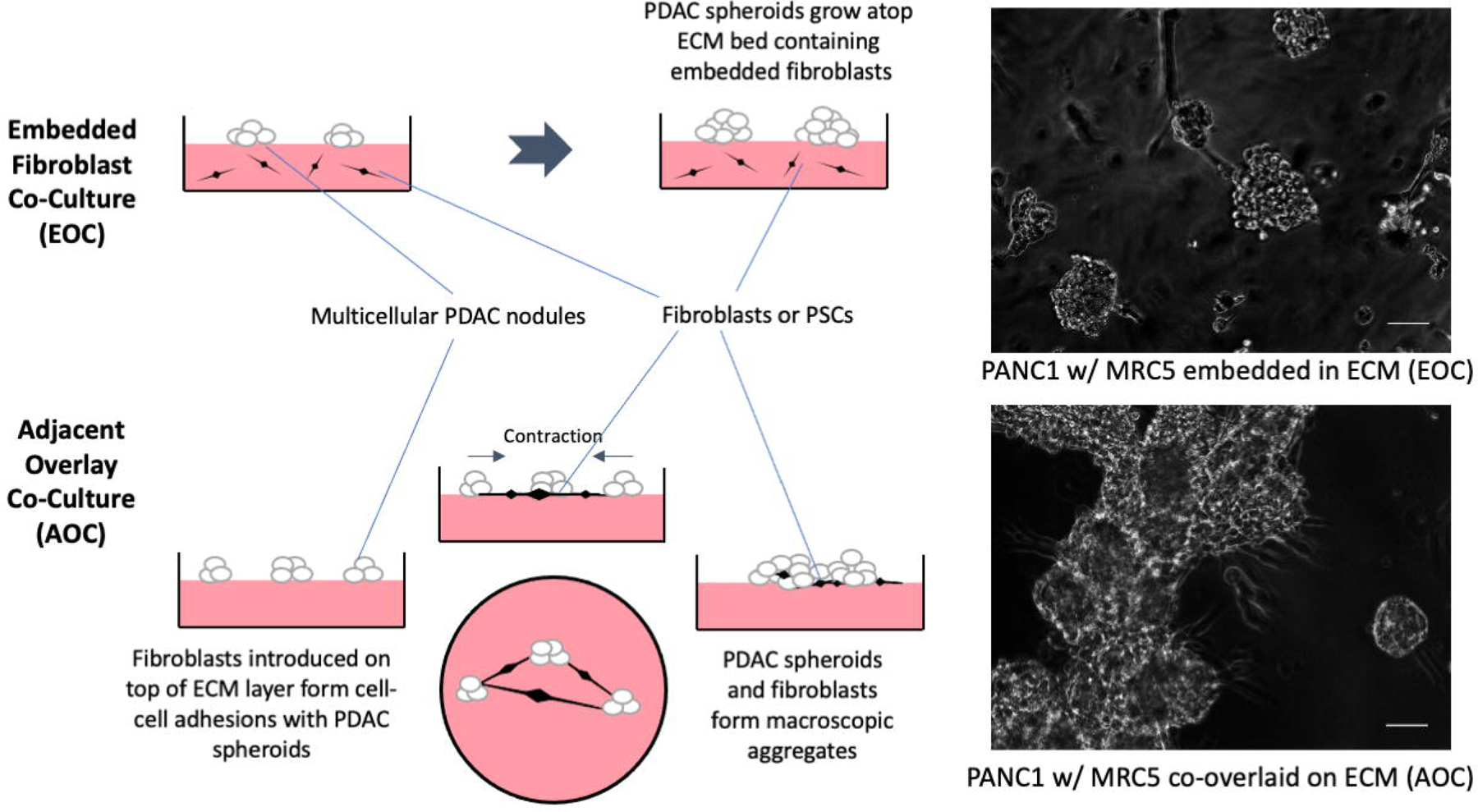
Overlay and adjacent tumor-fibroblast co-culture geometries. Left: Schematic diagrams of overlay and embedded co-culturing methods. In the embedded fibroblast with overlaid spheroid co-culture (EOC) fibroblasts are embedded in the underlying ECM layer and cannot make physical contact with PDAC spheroids overlayed above. In the adjacent fibroblast-spheroid overlay co-cuture (AOC) PDAC spheroids are grown on a Matrigel bed and fibroblasts are then introduced triggering contractile behaviour. Right: 10x phase contrast image data of each co-culture platform. The EOC image is taken after seven days of spheroid formation and the AOC image is taken 48 hours after fibroblasts have been introduced. Scale bars = 200μm

Co-cultures of adherent PDAC (PANC1) 3D nodules and fibroblastic cells (MRC5 or PSC) were initially carried out in two different geometries (**Figure 1**): 1) an embedded fibroblast/PDAC overlay co-culture (EOC) in which the fibroblastic cells are embedded in ECM but cancer cells are overlaid on the ECM surface, and 2) an adjacent overlay co-culture (AOC) in which both 3D PDAC nodules are formed on the spheroid surface and fibroblast are also added in the same plane. It has been shown that intercellular interactions between tumor cells and stroma are mediated by a combination of paracrine signaling and biophysical juxtacrine interactions which involve direct physical contact between tumor and stromal cells. When fibroblasts are suspended within the ECM bed and PANC-1 spheroids are grown on top (EOC), only paracrine signaling is able to occur by secretion of growth factors and cytokines that can diffuse through ECM. In EOC cultures the PANC1 nodule growth behavior is qualitatively similar to homotypic 3D cultures which have been extensively characterized in previous reports.^34–36^ Size distributions for PANC1 homotypic ECM overlay, PSC EOCs and MRC5 EOCs are shown in Supplemental Figure 1.

When fibroblasts are introduced on top of the ECM bed upon which adherent PANC-1 spheroids have been grown, both paracrine and juxtacrine interactions may occur. In this scenario, PDAC and fibroblastic cells rapidly co-migrate and form heterotypic adhesions. Once attached, the fibroblasts contract and pull the largely microscopic tumor spheroids into a macroscopic mass of mingled stroma and cancer cells near the center of the culture vessel. The 2D spatial distributions of nodules atop ECM bed after 48 hours in PANC-1 spheroid-fibroblast co-cultures show fewer and larger objects and with large swaths of empty ECM surface compared to PDAC spheroid homo-cultures. This is seen clearly in the PANC1-MRC5 AOC co-cultures where after 48 hours wells are dominated by single very large co-mingled nodule (**Supplemental Movie 1**). This data suggests that, at least in this *in vitro* co-culture geometry, interactions between PANC1 and fibroblasts are mediated by juxtacrine communication. This qualitative change in growth behavior prompted further image analysis to quantify changes in spheroid/fibroblast co-migration and contractile motion using the AOC model, which is the major focus of this study.

Analysis of time lapse image data from AOC cultures reveals contrasting fibroblast contractility observed in co-cultures with MRC5 versus those with PSCs (**Figure 2**). Here we observed that over the course of 48 hours, PANC1 cells grown in homoculture remain distributed across the ECM layer with small changes in position, while nodule-area weighted histograms confirmed an expected shift in size due to normal cell proliferation (Figure 2B). Distributions of nodule centroid positions similarly show there are fewer nodules but they remain distributed relatively uniformly across the well (Figure 2B). On the other hand, in MRC5 AOCs aggregation occurred rapidly, reslting in a large mass of PANC1 3D nodules enmeshed in fibroblasts. Area-weighted histograms highlight quantify aggregation with a dramatic shift towards few very large nodules over time. PSC AOC image data revealed signs of aggregation, however nodule size and centroid distribution data was less pronounced. Area-weighted histograms suggest a shift towards larger objects that isn’t particularly distinct from PANC1 homocultures, and centroid distibutions only hint at mild nodule aggregation with a slight increase in centroid-free space in the field of view. This contrast in contractility is consistent with previous characterization of co-cultures of these two cell types by us, suggesting more myofibroblastic CAF (MyCAF)-like phenotypic traits in MRC5 cells compared to the PSCs.^36^

**Figure 2:**
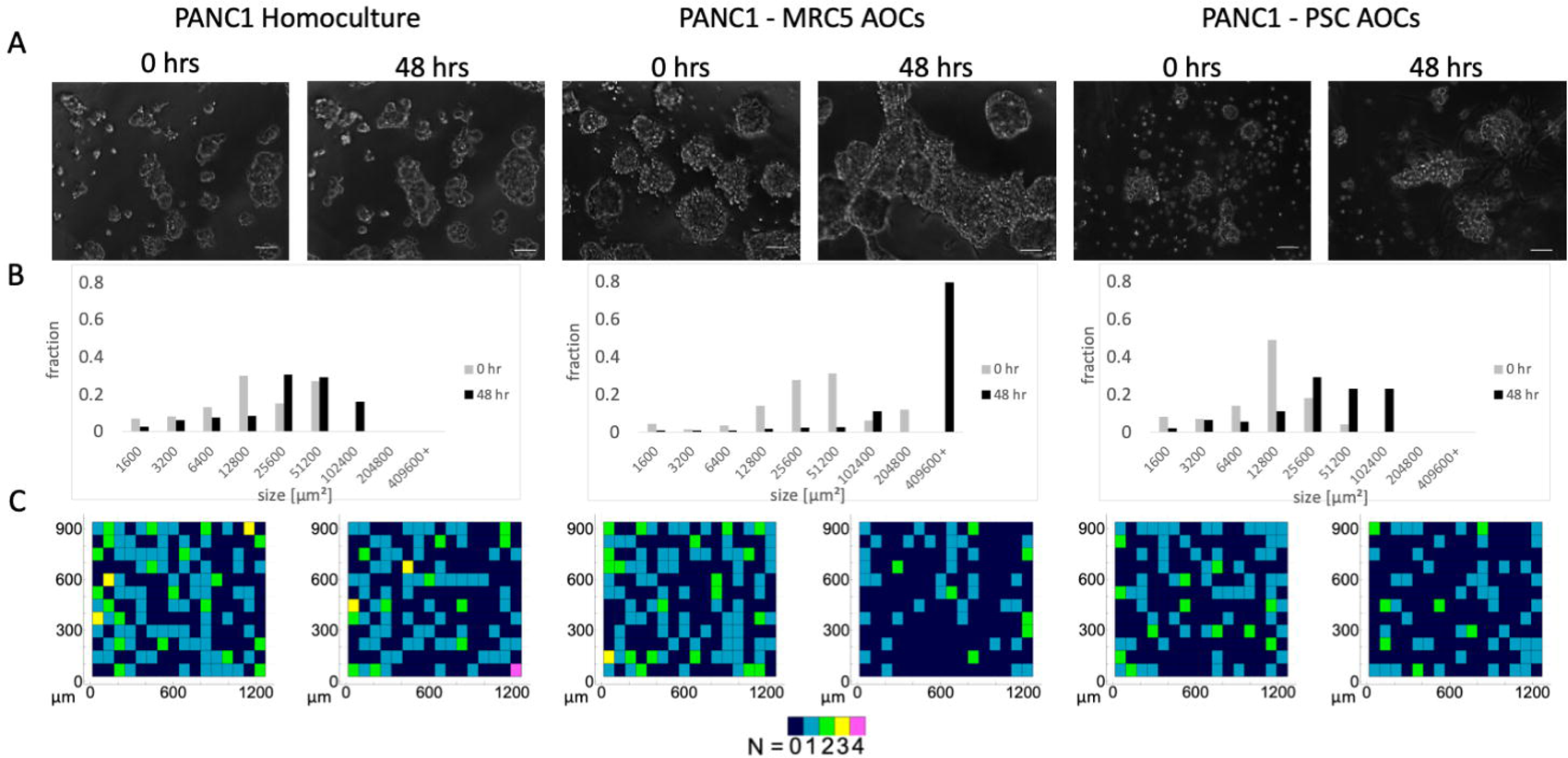
Contrasting growth behavior of homocultures and co-cultures of PDAC cells with fibroblasts or pancreatic stellate cells. **A)** Representative 10x phase contrast images from time lapse sequences of AOCs immediately (within ∼30 minutes) after introduction of fibroblasts and 48 hours later. Scale bars = 200 μm. **B**) Total area weighted histograms of nodule size distribution before and after initiation of fibroblast contractility. MyCAF-like MRC5 AOCs show a dramatic shift towards few very large objects, while PANC1 grown in homoculture and iCAF-like PSC data show only a subtle shift towards larger objects. **C)** Centroid distributions of nodules in each AOC at 0 and 48 hours. After 48 hours the MRC5 AOCs are spatially well separated on the ECM surface.

### 3.2 PIV analysis of PDAC homocultures and fibroblast co-cultures

PIV analysis was conducted to further analyze speed and patterns of migration and contractile motion occurring in PANC1 AOCs and PANC1 homocultures (Figure 3). This method involves quantification of shifts in object positions in segmented timelapse image sequences to generate a 2D map of velocity vectors at each time point (Supplemental Figure 2). From these velocity maps we calculate average velocity overall space in each each field as shown in Figure 3, where the error bars on each time point represent standard error in time resolved velocity measurements over all replicate movie sequences. The velocity profile for PANC-1 overlay homocultures is essentially flat (Figure 3A). In PSC AOCs (Figure 3B) there is some evidence of reproducible acceleration and deceleration, while in the MRC5 AOCs (Figure 3C) is there a well-defined and reproducible velocity profile that correlates with epithelial fibroblast co-migation, and fibroblast contraction events (Supplemental Figure 3). In the first 12 hours, a linear fit to the speed data shows acceraltion of 0.365 +/-0.019 μm/h^2^ with adjusted R-squared of 0.94. The deceleration phase afte 12 hours, corresponding to ongoing fibroblast contraction fits to an exponential decay in speed of the form *v* = *v*_0_ + *Ae^−t/τ^* with *v_0_* = 4.10 +/-0.034 μm/h^2^, *A* = 5.05 +/-0.27 μm/h^2^ and *τ* = 22.3 +/-1.3 hours, and adjusted R-squared of 0.85. The distinct velocity profile of AOC cultues contrasts with EOC experiments in which growth behavior of tumor nodules on the ECM surface that are not in direct contact with fibroblastic cells is qualitatively the same as in homocultures. Overall these experiments indicate the need for fibroblasts to have direct cell-cell juxtacrine interactions with cancer cells to induce dramatic contractile behavior, and that these interactions are most pronounced in the fibroblastic cells shown previously to exhibit a more myofibroblast-like phenotype.

**Figure 3:**
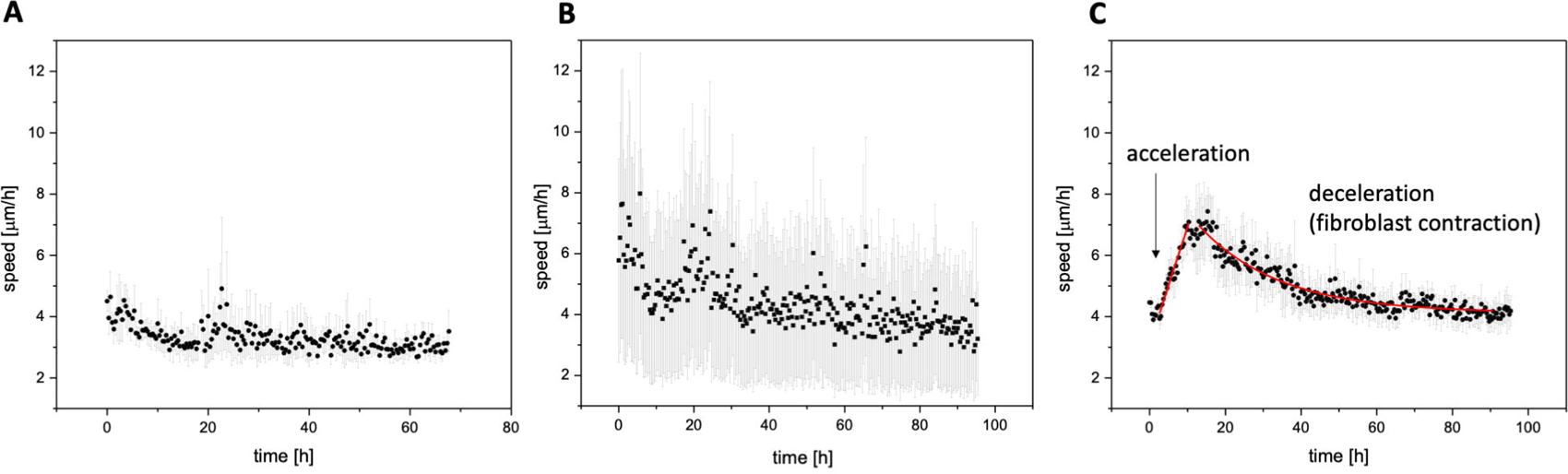
Particle image velocimetry analysis. Time-dependent velocimetry plots for PANC1 homocultures (**A**), PANC1 + PSC AOC (**B**) and PANC1 + MRC 5 AOC (**C**). Each data point shows mean velocity at each time point from 3 experiments and error bars indicate standard error. In homocultures the velocity profile is relatively flat while time dependent acceleration is evident in the two cultures. In the MRC5 AOCs velocity changes map to co-migration and fibroblast contraction events (further detailed in Supplemental Figure X) with linear increase in speed in the first 12 hours where in the first 12 hours and exponential decay after 12 hours (curve fits shown in red). The linear fit to the speed data shows initial acceraltion of 0.365 +/-0.019 μm/h^2^ with adjusted R-squared of 0.94. The deceleration phase afte 12 hours, corresponds to ongoing fibroblast contraction fits to an exponential decay in speed of the form *v* = *v*_0_ + *Ae^−t/τ^* with *v_0_* = 4.10 +/-0.034 μm/h^2^, *A* = 5.05 +/-0.27 μm/h^2^ and *τ* = 22.3 +/-1.3 hours, and adjusted R-squared of 0.85.

### 3.3 Altered behavior of adjacent overlay cultures of chemoresistant PDAC cells

To evaluate how interactions between PDAC and stromal cells is altered due to acquisition of chemoresistance we evaluated co-cultures of a chemoresistant subline of PANC1, PANC1-OR with previously reported resistance to oxaliplatin and gemcitabine chemotherapy, and which exhibit increased mesenchymal phenotypic traits and invasive behavior. ^22^ Here, following the same imaging and PIV analysis protocols described above, we examine contrasting growth behavior in PANC1 and PANC1OR AOC cultures.

**Figure 4:**
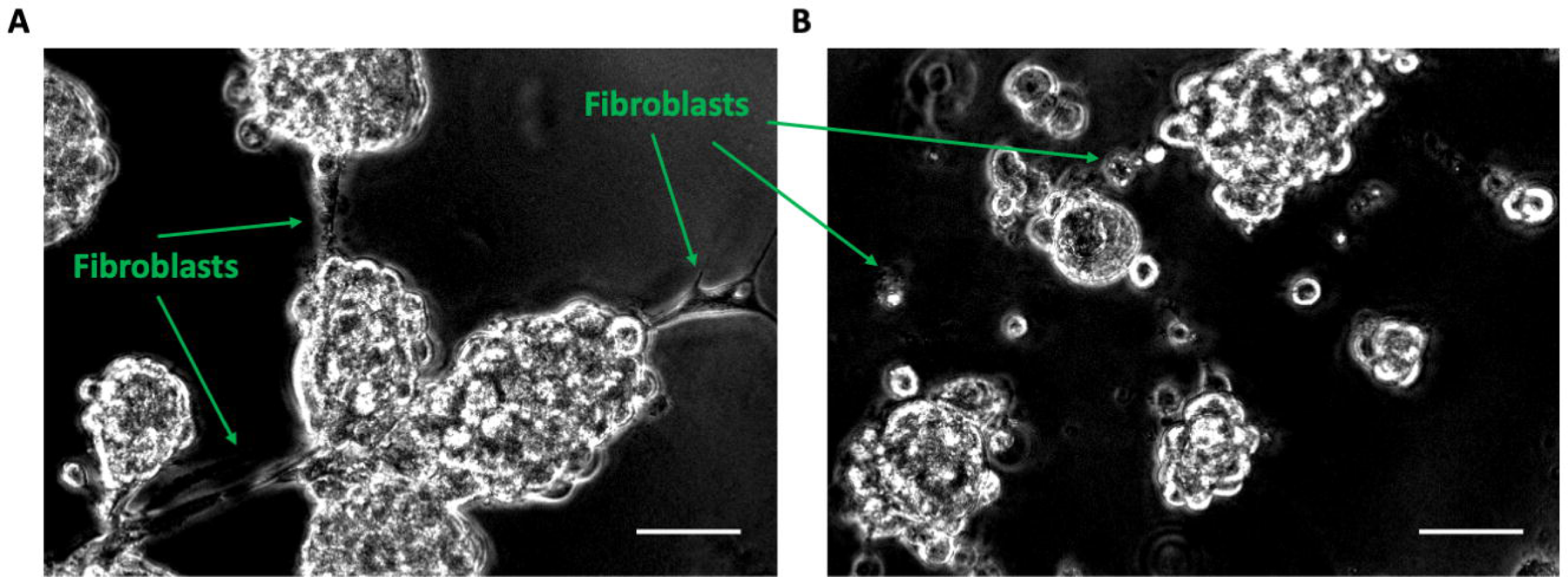
Representative images showing contrasting behavior of PANC1 versus PANC1-OR and MRC5 AOCs. **A)** 10x phase contrast image of a PANC1 with MRC5 AOC 12 hours after introduction of fibroblasts. Here fibroblasts make physical contact with spheroids leading to subsequent contraction. **B)** 10x phase contrast image of a PANC1-OR with MRC5 AOC 12 hours after addition of fibrobalsts in overlay cultures.

When grown on ECM beds, drug resistant PANC1-OR cells do form multicellular 3D nodules, but when MRC5 cells are introduced, there is a marked difference in the interaction with stromal cells when compared with drug-naïve PANC1 cells (**Figure 4**). Twelve hours after fibroblasts were introduced in PANC1 AOCs we observe fibroblasts stretched out and making physical contact with PANC1 nodules as they begin to merge into compact aggregates. (Figure 4A). In contrast, in PANC1-OR AOCs there are few extended fibroblasts and very few direct adhesions between PDAC nodules and fibroblasts (Figure 4B).

**Figure 5:**
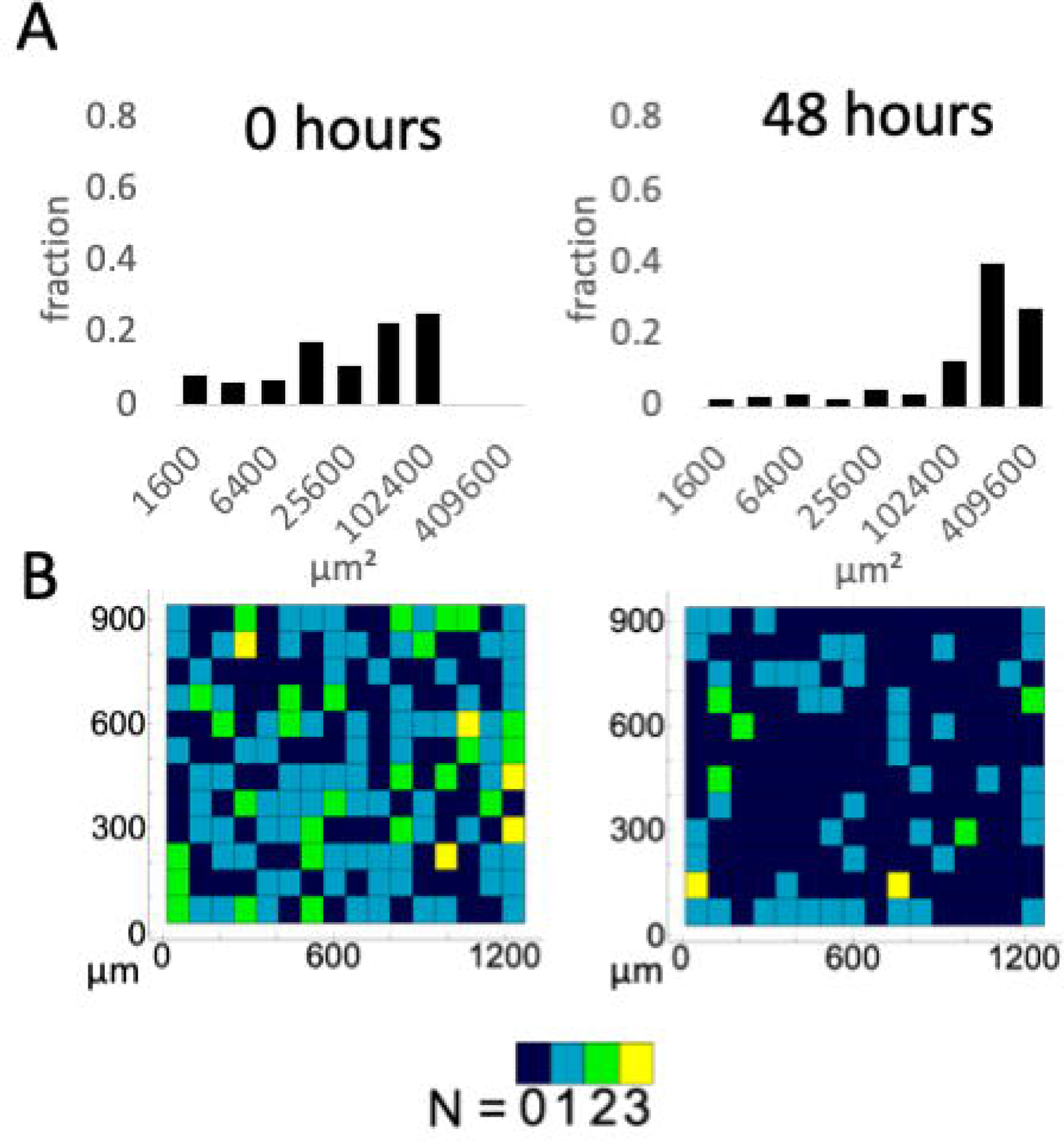
Analysis of size distributions over time for PANC1 versus PANC1-OR AOCs. **A)** Area weighted AOC nodule size histograms at 0 and 48 hours. PANC1-OR + MRC5 AOCs shift towards fewer and larger objects, though to a lesser extent than corresponding the PANC1 + MRC5 AOCs. **B)** Spatial distibutions at 0 and 48 hours after stromal cell overlay also show lower clustering relative to PANC1 + MRC5 AOCs shown in Figure 2.

Visualizing quantitative aggregation data, area-weighted nodule size histograms shows a decrease in aggregation in PANC1-OR AOCs (**Figure 5**) relative to PANC1 (non drug resistant) MRC5 AOCs shown in Figure 2. As seen in **Supplemental Movie 2**, the formation of adhesions between epithelial and stromal cells is less frequent than in co-cultures of the parental line and unaggregated cell clusters that persist through the duration of the experiment.

**Figure 6:**
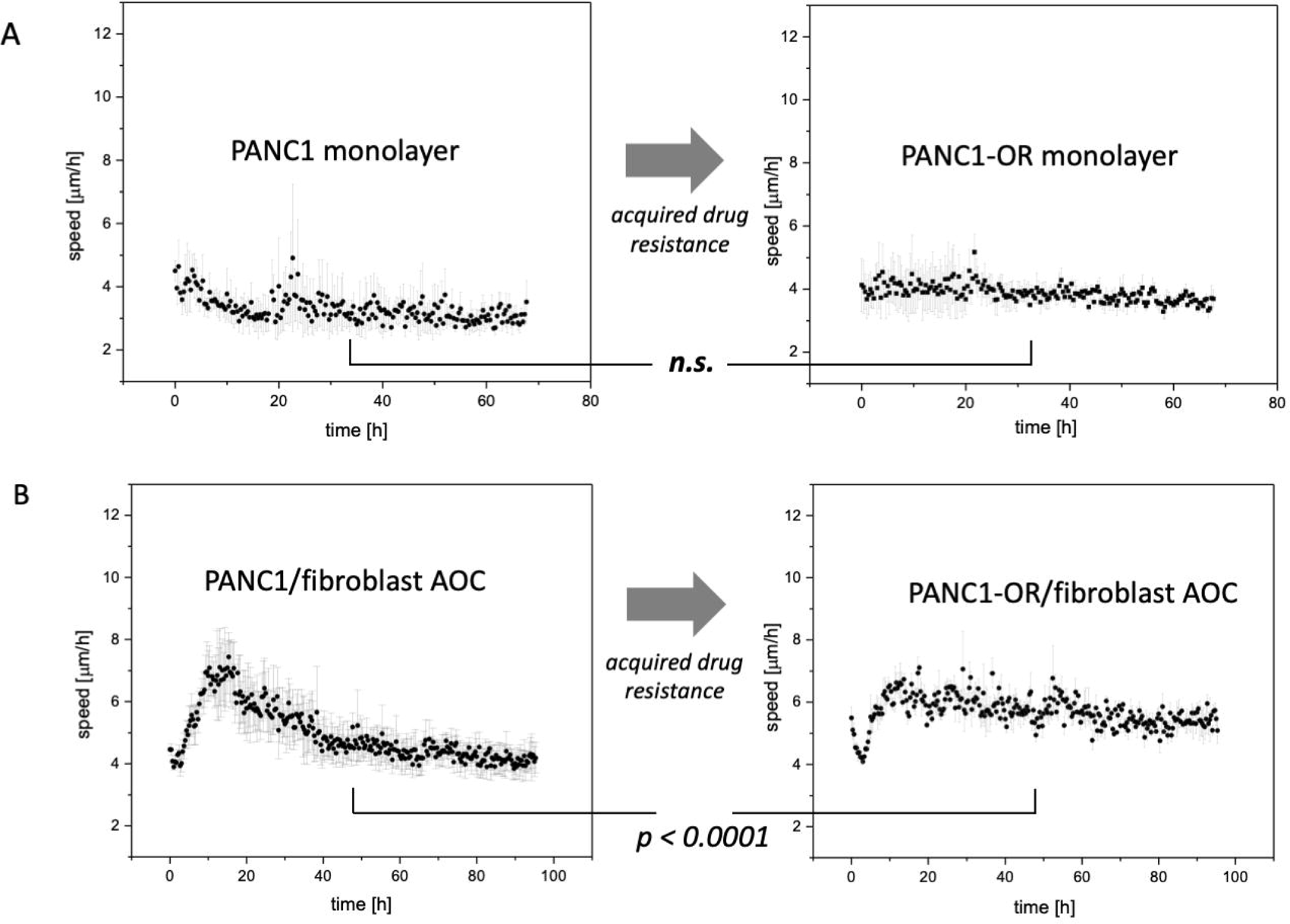
Comparative velocimetry analysis for PANC1 versus PANC1-OR AOCs. **A)** Comparison of velocimetry profiles for homocultures of PANC1 versus PANC1-OR. In both cases the overall time-dependent trend is flat with no significant change (p = 0.41 from analysis of sum of chi-squared weighted by standard error at each time point) with the acquisition of drug resistance. **B)** Comparison of mean speed in PANC1 vs PANC1-OR AOCs shows a highly signicant change. The well-defined acceleration phases of the PANC1 + MRC5 AOCs is no longer present when PANC1 are substituted with PANC1-OR. Chi-squared analysis shows the two curves different significantly with *p* < 0.001.

Comparison of velocimetry profiles obtained from sequenves of PANC1-OR and PANC1 co-cultures reveals marked change in behavior concomitant with acquisition of drug resistance (**Figure 6**). While chi-squared analysis comparing PANC1 and PANC1OR homoculture velocimetry plots (Figure 6A) shows no significant change (overall *p-value* of 0.41) the co-culture velocity curve (Figure 6B) changes significantly (*p <* 0.0001). The linear acceleration and exponential deceleration phases characterized above for PANC1 + MRC5 AOCs are less well defined the co-culture velocity profiles, which are generally noisier, and with higher mean speed over the duration of the experiment. This analysis is consistent with the qualitative observations that drug resistant cells are more motile and less likely to form cell-cell adhesions.

**Figure 7:**
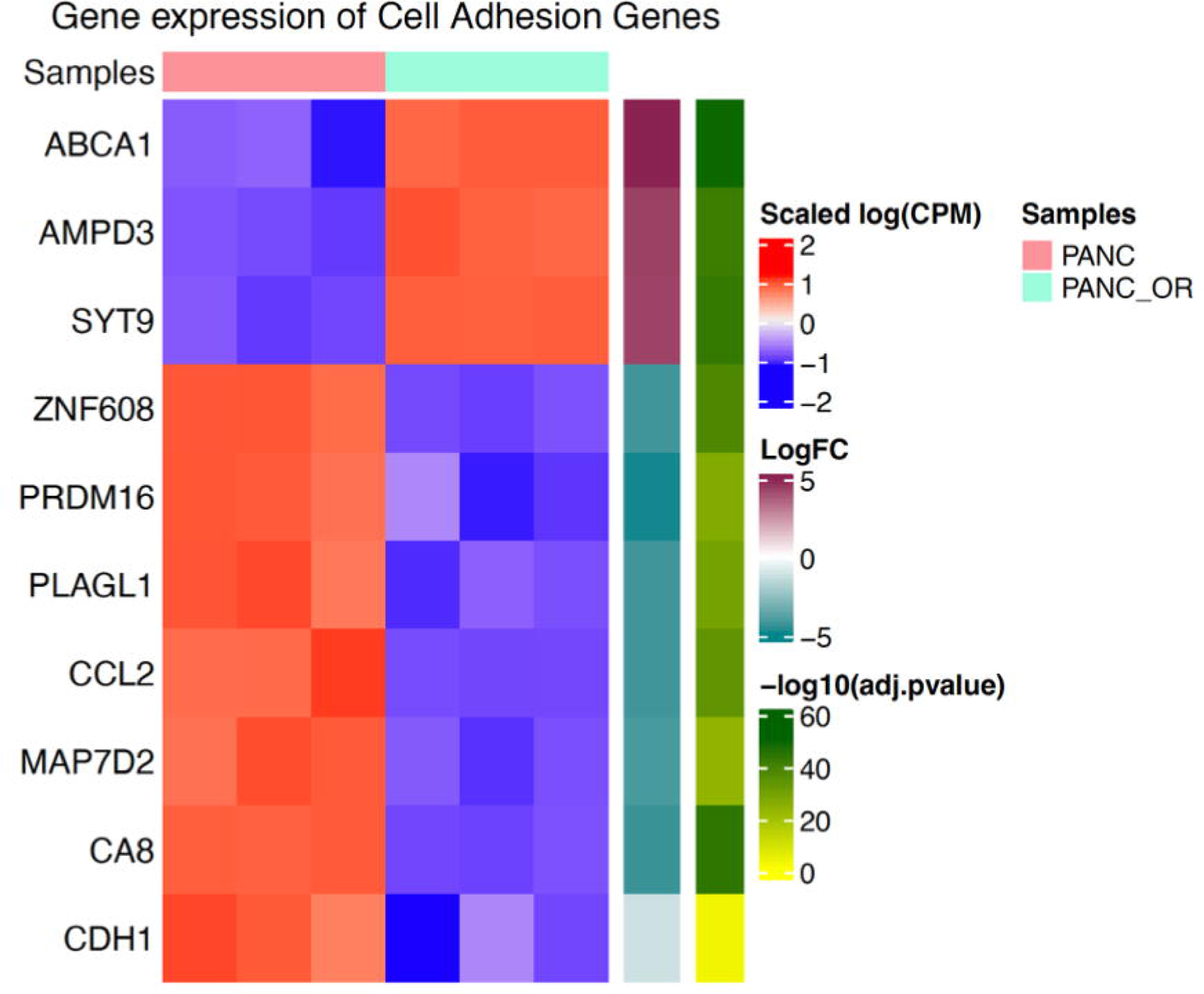
Heat map showing differential expression of genes relevant to cell-cell adhesion as determined from mRNA sequencing. All genes included were differentially expressed with *p* < 0.001. More comprehensive analysis of differential gene expression is shown in Supplemental Figures 3 through 7 and Supplemental Tables 1 to 3.

### 3.4 Analysis of differential gene expression in chemoresistant relative to chemosensitive PDAC cells

To examine changes in gene expression concomitant with acquisition of drug resistance that could account for suppression of adhesions to stromal cells observed above, we conducted mRNA sequencing and bioinformatics analysis to identify differentially expressed genes in PANC1-OR relative to PANC1. Extensive RNAseq analysis is shown **in Supplemental Figures 4 through 7** and **Supplemental Tables 1 to 3**. The gene ontology analysis for differential expression of gene familes related to biological processes, molecular functions, and cellular components in Supplemental Figures 3 through 5, respectively shows that PANC1-OR exhibit upregulation of genes related to cellular motility/migration, developmental and morphogenic processes, and decreased cell-cell adhesions. The overall pattern is generally consistent with increased EMT, though with some exceptions. The downregulation of CDH1 (log_2_(FC) = -1.2, *p* = 0.001), which codes for E-cadherin, is consistent with loss of epithelial characteristics. Other less obvious changes fit with this trend. For example, expression of KLF8, which has been shown to promote EMT by upregulating vimentin expression and downregulating E-cadherin,^37^ was increased in PANC1-OR (log_2_(FC) = +5.15, *p* = 8.09×10^-18^). On the other hand SNAI3 expression was decreased in Panc1OR, and SNAI1 expression was not significantly changed. Focusing on differential expression of genes potentially relevant to the observations here we found broadly that there was significantly decreased expression of genes which code for cell adhesion molecules in PANC1-OR (Figure 7). Immunofluorescene imaging of fixed cultures (**Supplemental Figure 8**) also shows loss of E-cadherin in PANC1-OR. Also, while the vimentin gene was not differentially expressed, the immunofluorescence imaging shows a clear pattern of cell elongation with increased cytoskeletal vimentin in PANC1-OR. Further phenotypic analysis of these drug resistant cells has been reported previously.^22^

## Conclusion

In this study we use a set of image analysis tools to contrast growth behavior in adjacent overlay PDAC 3D co-cultures. The analysis performed in this study points to the important role of direct biophysical interaction requiring cell-cell contacts between epithelial and stromal cells in PDAC. Only when fibroblasts are free to physically attach to PDAC nodules (as in the AOC model), do we see strong evidence of direct cell-cell contacts which mediate fibroblast contreactility. This growth behavior drives toward densely compressed aggregates of intermingled tumor and stromal cells. This system could be further leveraged to model desmoplastic PDAC tissues and study how compressive stress in fibrotic stroma impacts upon disease progression in a controlled laboratory model.

A major goal of this study was to examine how these biophysical interactions between epithelial and adjacent stromal cells become altered in drug resistant PDAC. We observe through imaging and image analysis that the physical contacts required for contractility and aggregation are suppressed in drug resistant as compared to drug naïve PDAC cells co-cultured with fibroblasts. This finding is supported by RNA sequencing and immunofluorescence imaging showing dwonregulation of genes required for cell-cell adhesions between tumor and stromal cells, which is consistent with increased EMT and invasive behavior in these cells. This is significant in view of evidence that these contractile interactions between epithelial PDAC and myofiobroblastic cancer associated fibroblast subpopulations act to constrain invasive progression of disease.^38^ On the other hand it is also important to note that heterophilic E-cadherin/N-cadherin adhesions between cancer cells and stromal fibroblasts have been shown to promote an invasion mechanism in which CAFs guide collective migration of epithelial cells with intact adherens junction.^39^ However, the drug resistant cells studied here, which have almost no expression of E-cadherin at the gene or protein level, would not be able to participate in this collective migration. Taken together, this suggests that single cell EMT-mediated invasion could be enabled by loss of adhesion to myofibroblastic CAFs which could otherwise contribute to mechanical confinement through exertion of contractile compressive stresses as seen in co-cultures with the parent PDAC line. In other words, the acquisition of drug resistance, which itself contributes to poor outcomes in PDAC,^20^ is found here to also contribute to inhibition of stromal interactions that could otherwise help constrain invasive PDAC progression.

In these imaging-based biophysical studies we found that MRC5 co-cultures were more chartacteristically myofibroblastic, producing profound contractile behavior and much greater shift in object size and clustering than PSC aggregates, with primary nodules encompassing the majority of cells in the well. Further MRC5 AOCs result in a nearly twice as strong contractile response as PSC AOCs. MyCAF interactions are known to be mediated by juxtacrine signaling, and the differential response between MRC5 and PSC AOCs suggests fibroblast contractility, and that the intercellular forces induced by stromal crosstalk, are mediated by juxtacrine interactions within the PDAC tumor microenvironment.

The macroscopic AOC aggregates cultured in this study may also be a useful tool to recapitulate desmoplasia characteristic of human PDAC in a versatile *in vitro* experimental platform. From the time evolution of contractile behavior we can glean insight into the mechanics of these tumor fibroblast interactions. Also, as a 3D culture methodology, the development of these highly fibrotic intermingled tumor and stroma tissue models could be used to study drug delivery in desmoplastic tumors,^40^ or detacted frm ECM for xenograft implantation or further downstream analysis.

## Data Availability

The datasets used and/or analysed during the current study available from the corresponding author on reasonable request.

## Acknowledgements

We gratefully acknowledge funding from NIH/NCI grants, R35CA232105 (FJS) and U54CA156734 (JPC and FJS), which supports the UMass Boston-Dana Farber/Harvard Cancer Center Partnership, and the University of Massachusetts Boston Healey Research Grant Program (JBD and JPC). We would also like to thank Fathi Mohamed for helping prepare samples for RNA sequencing.

## Author Contributions Statement

E.S performed experiments and prepared the initial manuscript draft. J.P.C. guided experimental design and revised manuscript text. F.J.S. and J.B.D. guided experimental design and revised manuscript text.

G.M.C. generated and characterized drug resistant cells. V.K. contributed to design of cell culture methods and data for Figure 1. M.L. carried out bioinformatics analysis under supervision of K.Z. Y.L. assisted with bioinformatics analysis and writing. M.L. and K.Z. also revised the manuscript text.

