## Supplemental Figures and Tables for "Drug resistant pancreatic cancer cells exhibit altered biophysical interactions with stromal fibroblasts in imaging studies of 3D co-culture models"

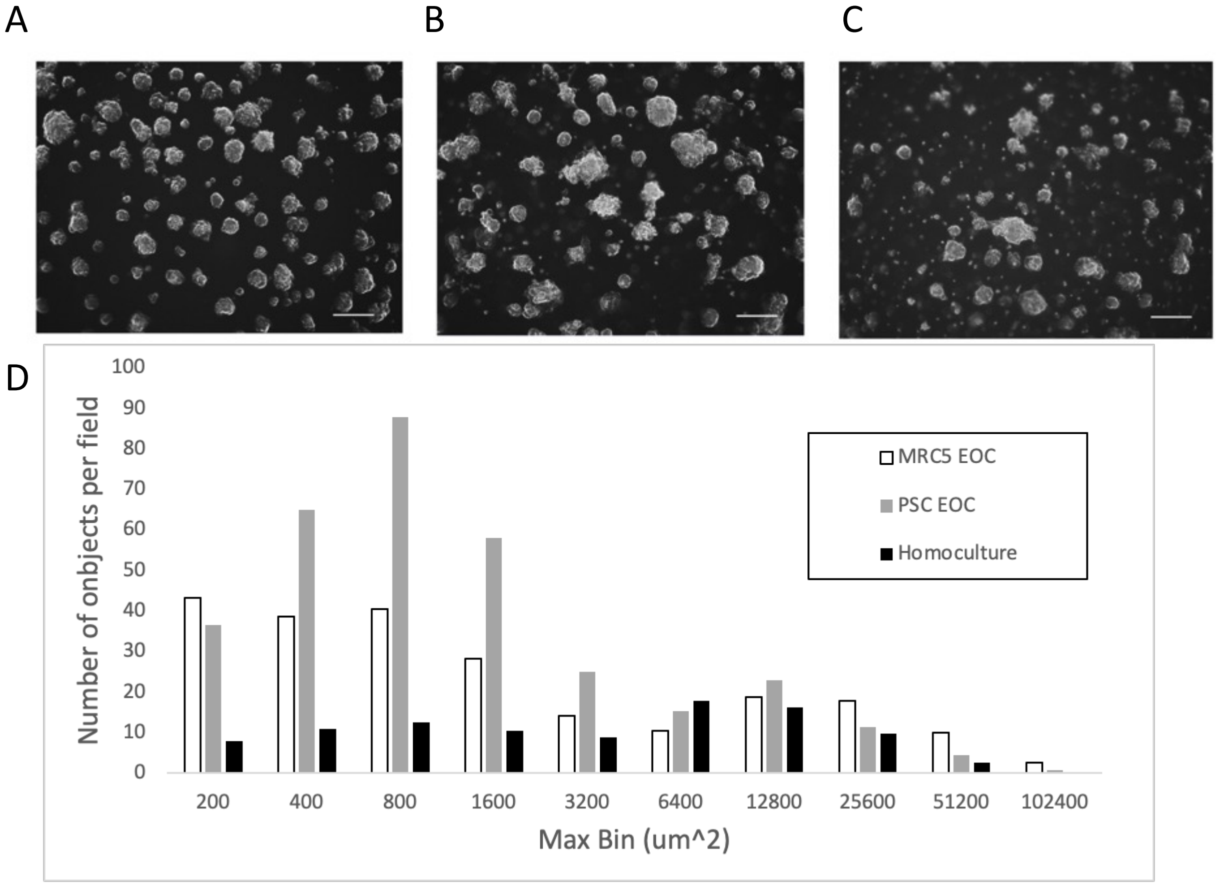

**Supplemental Figure 1:** Representative 5X darkfield snapshots of **A**) PANC1 3D Matrigel overlay homocultures, B) PANC1/MRC5 EOCs, and C) PANC1/PSC EOCs, imaged 7 days after plating, and D) a histogram showing the size distribution for each condition. Size distributions are scaled by the number of fields imaged (8 to 12 for each condition). The distribution of larger spheroids is similar in each condition with a slightly higher proportion of large nodules in MRC5 EOCs. Higher numbers of small objects in co-cultures relative to PANC1 homoculture is largely due to the fibroblastic cells themselves. None of the EOC conditions exhibit the dramatic aggregation and contractile behavior which is the major focus of this study.

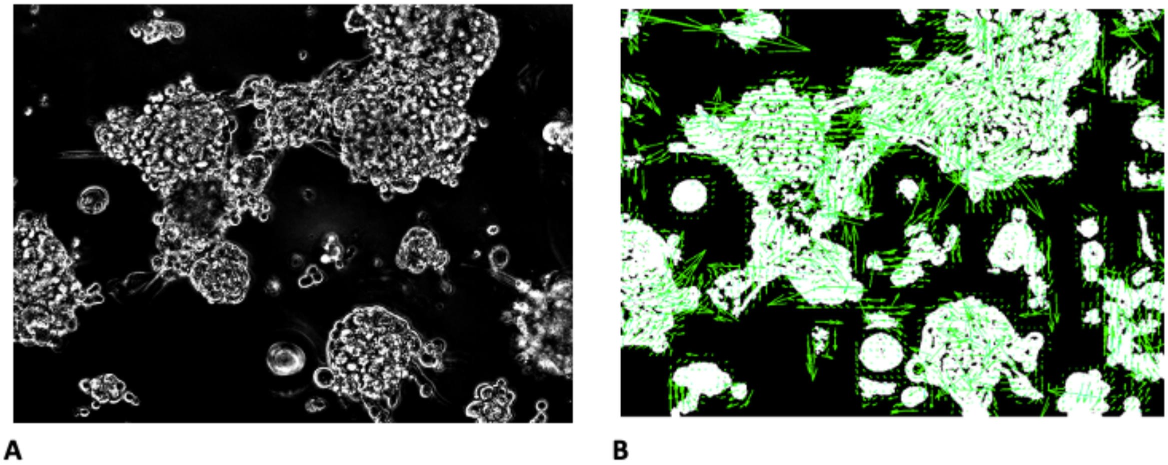

**Supplemental Figure 2: A**) A representative 10 X phase contrast image at timepoint of peak nodule speed during in-progress nodule aggregation in a PANC1-MRC5 AOC model. **B**) The corresponding segmented binary image with overlaid velocity vector map generated by PIVlab. This is representative of the velocity vector maps used throughout this study to calculate velocity and acceleration for various times and cell culture conditions.

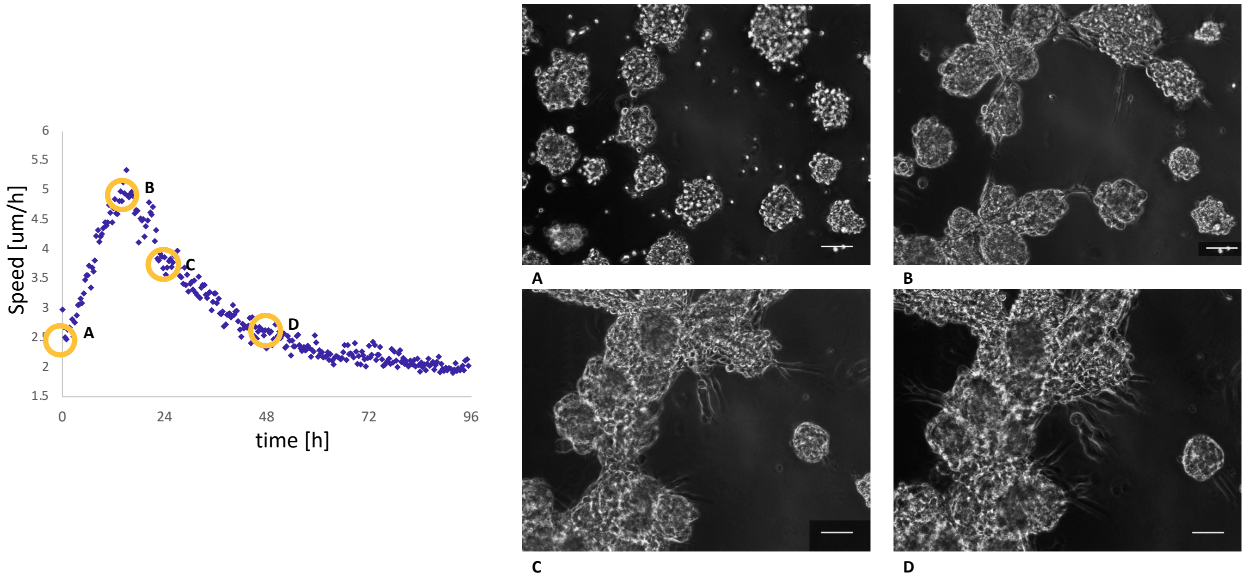

**Supplemental Figure 3:** A representative velocimetry profile for a PANC1/MRC5 AOC showing characteristic acceleration and velocity relaxation phases correlated with aggregation events from indicated timelapse frames (right). From [A] to [B] speed increase is linear, followed by exponentially decreasing speed beyond 12 hours, correlated with continued fibroblast contraction [C] and [D]. Large intermingled tumor spheroid/fibroblast aggregates continue to pull together with exponentially decreasing speed up to 48 hours [D] and beyond. Scale bars = 200 μm.

A

B

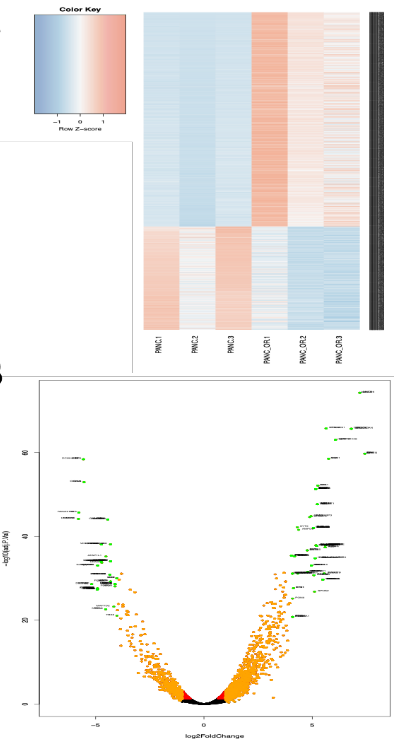

C

| Ensemble | entrez | Symbol | logFC | PValue | biotype |
|---|---|---|---|---|---|
| ENSG00000187957 | 92737 | DNER | 7.19827452 | 5.61E-75 | protein_coding |
| ENSG00000074527 | 59277 | NTN4 | 5.63762909 | 1.76E-66 | protein_coding |
| ENSG00000152377 | 6695 | SPOCK1 | 6.80354013 | 2.15E-66 | protein_coding |
| ENSG00000176658 | 4642 | MYO1D | 6.0726316 | 9.51E-64 | protein_coding |
| ENSG00000124882 | 2069 | EREG | 7.43096366 | 1.81E-60 | protein_coding |
| ENSG00000118515 | 6446 | SGK1 | 5.75541938 | 3.07E-59 | protein_coding |
| ENSG00000096696 | 1832 | DSP | -5.5642645 | 3.92E-59 | protein_coding |
| ENSG00000272398 | 100133941 | CD24 | -5.5347115 | 1.14E-53 | protein_coding |
| ENSG00000154027 | 26289 | AK5 | 5.25850443 | 7.75E-53 | protein_coding |
| ENSG00000152661 | 2697 | GJA1 | 5.16339134 | 5.13E-52 | protein_coding |
| ENSG00000165029 | 19 | ABCA1 | 5.23907083 | 2.23E-48 | protein_coding |
| ENSG00000132688 | 10763 | NES | -5.7747746 | 2.07E-46 | protein_coding |
| ENSG00000152689 | 25780 | RASGRP3 | 4.94478685 | 1.57E-45 | protein_coding |
| ENSG00000198879 | 57713 | SFMBT2 | 4.88121324 | 2.52E-45 | protein_coding |
| ENSG00000182168 | 8633 | UNC5C | -5.7921774 | 6.75E-45 | protein_coding |
| ENSG00000178538 | 767 | CA8 | -4.4395955 | 8.90E-45 | protein_coding |
| ENSG00000170743 | 143425 | SYT9 | 4.31264586 | 6.34E-43 | protein_coding |
| ENSG00000198947 | 1756 | DMD | 5.05455035 | 9.49E-43 | protein_coding |
| ENSG00000133805 | 272 | AMPD3 | 4.37028166 | 2.75E-42 | protein_coding |
| ENSG00000106123 | 20511 | EPHB6 | 6.09932452 | 2.26E-40 | protein_coding |
| ENSG00000168916 | 57507 | ZNF608 | -4.3146449 | 7.60E-39 | protein_coding |
| ENSG00000165810 | 153579 | BTNL9 | -4.7382146 | 8.00E-39 | protein_coding |
| ENSG00000121966 | 7852 | CXCR4 | 5.62514102 | 1.26E-38 | protein_coding |
| ENSG00000170381 | 9723 | SEMA3E | 5.17631924 | 1.38E-38 | protein_coding |
| ENSG00000067798 | 89795 | NAV3 | 5.202662 | 1.91E-38 | protein_coding |
| ENSG00000005981 | 51666 | ASB4 | 5.59619762 | 3.96E-38 | protein_coding |
| ENSG00000067715 | 6857 | SYT1 | 4.76831936 | 2.17E-37 | protein_coding |
| ENSG00000198682 | 9060 | PAPSS2 | 4.03936241 | 4.12E-36 | protein_coding |
| ENSG00000163347 | 9076 | CLDN1 | 4.18248298 | 5.40E-36 | protein_coding |

**Supplemental Figure 4:** Analysis differentially expressed genes (DEGs) obtained from RNAseq of PANC1 and PANC1OR cell cultures calculated using edgeR package. (A) Raw counts for all DEGs from triplicate independent samples of PANC1 and PANC1 OR cell cultures. There are 1342 protein coding genes with abs(log2FC)> 2 and p.value < 0.01. The heat map shows the separation of upregulated and downregulated genes in two groups. (B) Volcano plot. The log2(FC) is on the x-axis and –log(p-value) on the y-axis. A positive log2(FC) shows the genes that are upregulated in PANC1_OR. The genes in green font are DEGs with abs(log2FC) > 4 and p-value < 0.001. (C) A table of the top DEGs (highest fold change, positive or negative) in PANC1-OR relative to PANC1.

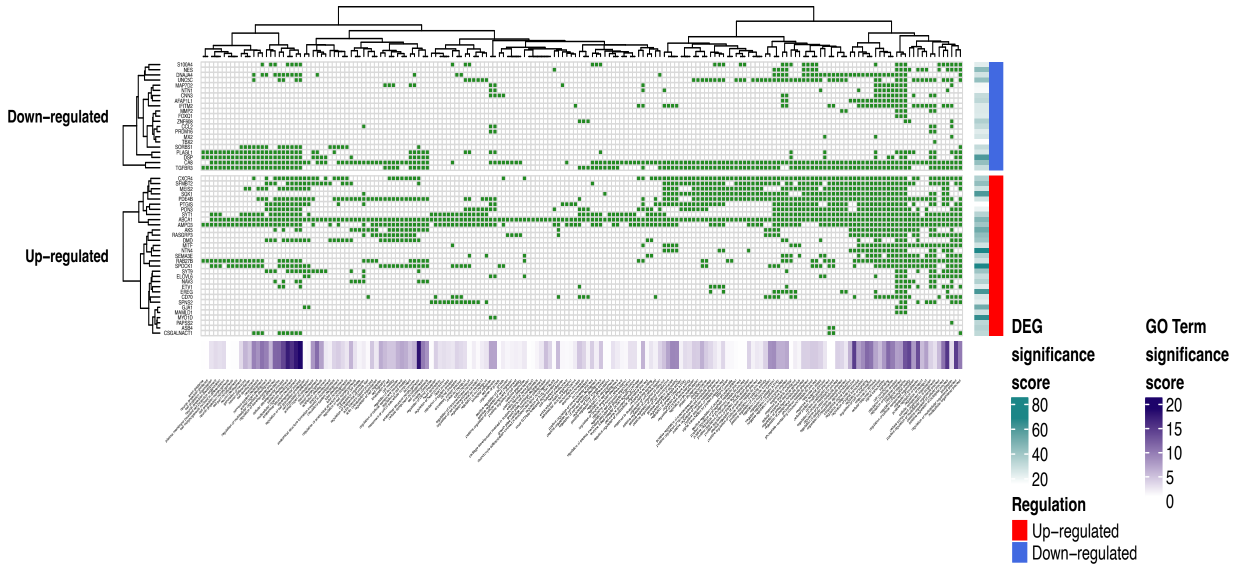

**Supplemental Figure 5:** Gene Ontology (GO) enrichment analysis for biological processes corresponding. This heat map plot of GO term annotations and DEGs corresponds to the list in Supplemental Table 1. Significant GO terms are on the rows and their associated gene targets from DEGs table are in the columns.

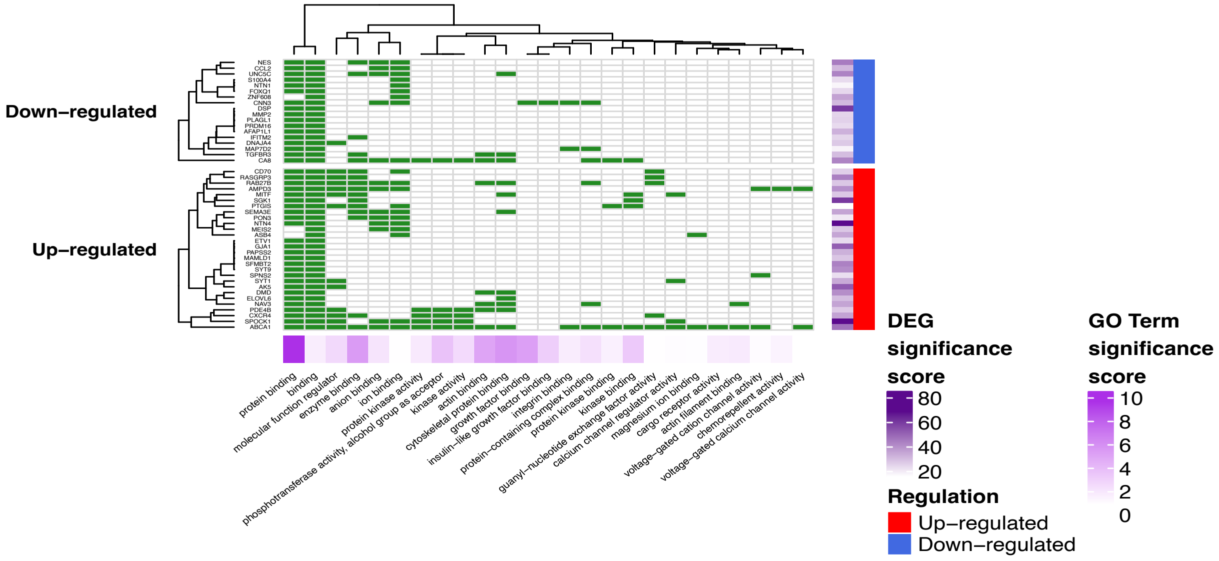

**Supplemental Figure 6:** Gene Ontology (GO) enrichment analysis for Molecular Function. This heat map plot of GO term annotations and DEGs corresponds to the list in Supplemental Table 2. Significant GO terms are on the rows and their associated gene targets from DEGs table are in the columns.

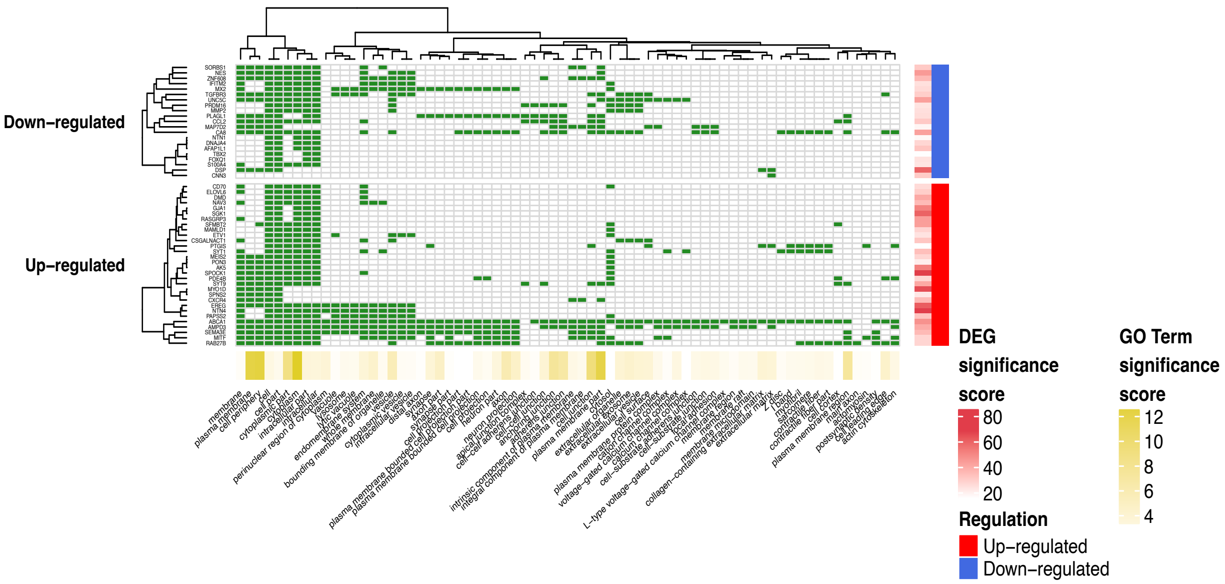

**Supplemental Figure 7:** Gene Ontology (GO) enrichment analysis for Cellular Components. This heat map plot of GO term annotations and DEGs corresponds to the list in Supplemental Table 3. Significant GO terms are on the rows and their associated gene targets from DEGs table are in the columns.

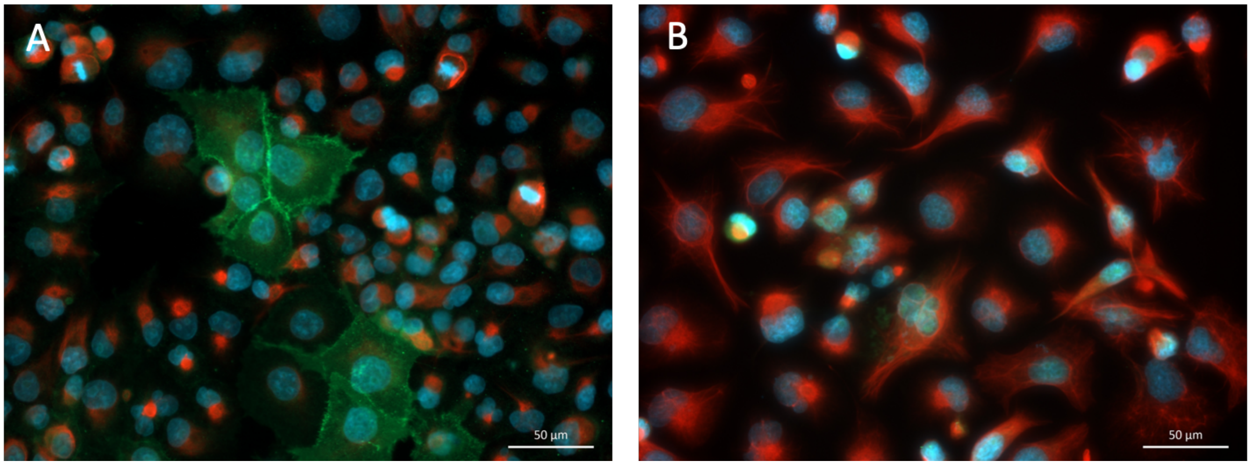

**Supplemental Figure 8:** Immunofluorescence imaging of (A) PANC1, and (B) PANC1-OR cells. E-cadherin staining is shown in green, vimentin in red, and DAPI (nuclei) in blue. Drug resistant cells exhibit decreased E-cadherin and increased cytoskeletal vimentin, consistent with increased EMT.

| source | term_name | term_id | adjusted_p_value | negative_log10_of_adjusted_p_value | term_size | query_size | intersection_size | effective_domain_size |
|---|---|---|---|---|---|---|---|---|
| GO:BP | developmental process | GO:0032502 | 4.27E-19 | 18.3693892 | 6205 | 1196 | 579 | 17622 |
| GO:BP | anatomical structure development | GO:0048856 | 2.00E-17 | 16.6979566 | 5813 | 1196 | 544 | 17622 |
| GO:BP | system development | GO:0048731 | 3.56E-17 | 16.4487681 | 4801 | 1196 | 468 | 17622 |
| GO:BP | anatomical structure morphogenesis | GO:0009653 | 8.31E-17 | 16.0801524 | 2631 | 1196 | 295 | 17622 |
| GO:BP | positive regulation of biological process | GO:0048518 | 2.39E-16 | 15.6207208 | 6039 | 1196 | 556 | 17622 |
| GO:BP | multicellular organism development | GO:0007275 | 1.19E-15 | 14.9238036 | 5339 | 1196 | 502 | 17622 |
| GO:BP | intracellular signal transduction | GO:0035556 | 4.42E-15 | 14.3542675 | 2751 | 1196 | 299 | 17622 |
| GO:BP | cellular developmental process | GO:0048869 | 8.21E-15 | 14.0858641 | 4306 | 1196 | 421 | 17622 |
| GO:BP | regulation of multicellular organismal process | GO:0051239 | 9.84E-15 | 14.0071509 | 3098 | 1196 | 326 | 17622 |
| GO:BP | biological regulation | GO:0065007 | 1.64E-14 | 13.786012 | 12195 | 1196 | 955 | 17622 |
| GO:BP | regulation of localization | GO:0032879 | 6.84E-14 | 13.1647627 | 2718 | 1196 | 292 | 17622 |
| GO:BP | regulation of biological quality | GO:0065008 | 1.38E-13 | 12.8606283 | 3864 | 1196 | 382 | 17622 |
| GO:BP | signaling | GO:0023052 | 3.79E-13 | 12.4216082 | 6457 | 1196 | 572 | 17622 |
| GO:BP | cell differentiation | GO:0030154 | 4.77E-13 | 12.3215209 | 4127 | 1196 | 400 | 17622 |
| GO:BP | regulation of signaling | GO:0023051 | 5.49E-13 | 12.2604894 | 3536 | 1196 | 354 | 17622 |
| GO:BP | tissue development | GO:0009888 | 1.06E-12 | 11.973431 | 1980 | 1196 | 226 | 17622 |
| GO:BP | positive regulation of cellular process | GO:0048522 | 2.50E-12 | 11.601531 | 5291 | 1196 | 484 | 17622 |
| GO:BP | regulation of cell communication | GO:0010646 | 2.65E-12 | 11.5770154 | 3508 | 1196 | 349 | 17622 |
| GO:BP | regulation of developmental process | GO:0050793 | 3.93E-12 | 11.405333 | 2615 | 1196 | 277 | 17622 |

**Supplemental Table 1:** Gene ontology enrichment analysis for Biological Processes.

Enrichment analysis for DEGs was run using https://david.ncifcrf.gov.

| source | term_name | term_id | adjusted_p_value | negative_log10_of_adjusted_p_value | term_size | query_size | intersection_size | effective_domain_size |
|---|---|---|---|---|---|---|---|---|
| source | term_name | term_id | adjusted_p_value | negative_log10_of_adjusted_p_value | term_size | query_size | intersection_size | effective_domain_size |
| GO:MF | protein binding | GO:0005515 | 7.10E-11 | 10.1490077 | 11632 | 1213 | 919 | 17516 |
| GO:MF | enzyme binding | GO:0019899 | 3.85E-07 | 6.41442604 | 2266 | 1213 | 229 | 17516 |
| GO:MF | cytoskeletal protein binding | GO:0008092 | 3.32957E-06 | 5.47761225 | 937 | 1213 | 112 | 17516 |
| GO:MF | actin binding | GO:0003779 | 1.62335E-05 | 4.78958836 | 422 | 1213 | 61 | 17516 |
| GO:MF | growth factor binding | GO:0019838 | 2.47527E-05 | 4.60637798 | 137 | 1213 | 29 | 17516 |
| GO:MF | kinase binding | GO:0019900 | 0.000103137 | 3.98658342 | 719 | 1213 | 87 | 17516 |
| GO:MF | molecular function regulator | GO:0098772 | 0.000336517 | 3.47299341 | 1864 | 1213 | 182 | 17516 |
| GO:MF | phosphotransferase activity, alcohol group as acceptor | GO:0016773 | 0.000342018 | 3.46595122 | 795 | 1213 | 92 | 17516 |
| GO:MF | insulin-like growth factor binding | GO:0005520 | 0.000595144 | 3.22537823 | 28 | 1213 | 11 | 17516 |
| GO:MF | protein-containing complex binding | GO:0044877 | 0.000606563 | 3.21712444 | 1070 | 1213 | 115 | 17516 |
| GO:MF | anion binding | GO:0043168 | 0.001188877 | 2.92486293 | 2780 | 1213 | 251 | 17516 |
| GO:MF | kinase activity | GO:0016301 | 0.002640153 | 2.57837096 | 892 | 1213 | 97 | 17516 |
| GO:MF | protein kinase binding | GO:0019901 | 0.004189067 | 2.37788264 | 639 | 1213 | 74 | 17516 |
| GO:MF | cargo receptor activity | GO:0038024 | 0.008270961 | 2.08244404 | 85 | 1213 | 18 | 17516 |
| GO:MF | integrin binding | GO:0005178 | 0.009194483 | 2.03647268 | 127 | 1213 | 23 | 17516 |
| GO:MF | actin filament binding | GO:0051015 | 0.011213849 | 1.95024531 | 192 | 1213 | 30 | 17516 |
| GO:MF | protein kinase activity | GO:0004672 | 0.012127334 | 1.91623465 | 659 | 1213 | 74 | 17516 |
| GO:MF | binding | GO:0005488 | 0.013569534 | 1.86743506 | 15096 | 1213 | 1091 | 17516 |
| GO:MF | chemorepellent activity | GO:0045499 | 0.01870878 | 1.72795453 | 26 | 1213 | 9 | 17516 |
| GO:MF | carbohydrate derivative binding | GO:0097367 | 0.02421438 | 1.61592664 | 2230 | 1213 | 200 | 17516 |

**Supplemental Table 2:** Gene ontology enrichment analysis for molecular function. Enrichment analysis for DEGs was run using https://david.ncifcrf.gov.

| source | term_name | term_id | adjusted_p_value | negative_log10_of_adjusted_p_value | term_size | query_size | intersection_size | effective_domain_size |
|---|---|---|---|---|---|---|---|---|
| GO:CC | cytoplasm | GO:0005737 | 2.39E-13 | 12.6212966 | 11436 | 1261 | 901 | 18745 |
| GO:CC | cell periphery | GO:0071944 | 5.83E-13 | 12.2342258 | 5586 | 1261 | 503 | 18745 |
| GO:CC | plasma membrane | GO:0005886 | 9.79E-13 | 12.0093333 | 5477 | 1261 | 494 | 18745 |
| GO:CC | plasma membrane part | GO:0044459 | 2.14E-12 | 11.6698985 | 2830 | 1261 | 291 | 18745 |
| GO:CC | cell junction | GO:0030054 | 1.94E-11 | 10.7124013 | 1280 | 1261 | 157 | 18745 |
| GO:CC | plasma membrane region | GO:0098590 | 6.37E-10 | 9.19608537 | 1144 | 1261 | 140 | 18745 |
| GO:CC | anchoring junction | GO:0070161 | 7.65E-09 | 8.11624833 | 552 | 1261 | 81 | 18745 |
| GO:CC | adherens junction | GO:0005912 | 3.01E-08 | 7.52202495 | 537 | 1261 | 78 | 18745 |
| GO:CC | cytoplasmic part | GO:0044444 | 4.20E-08 | 7.37703651 | 9606 | 1261 | 755 | 18745 |
| GO:CC | axon | GO:0030424 | 7.81E-07 | 6.10761982 | 596 | 1261 | 80 | 18745 |
| GO:CC | synapse | GO:0045202 | 1.4756E-06 | 5.8310354 | 1034 | 1261 | 119 | 18745 |
| GO:CC | vesicle | GO:0031982 | 1.4826E-06 | 5.82896534 | 3811 | 1261 | 339 | 18745 |
| GO:CC | neuron projection | GO:0043005 | 2.4895E-06 | 5.60388372 | 1300 | 1261 | 141 | 18745 |
| GO:CC | neuron part | GO:0097458 | 5.2554E-06 | 5.27939647 | 1666 | 1261 | 170 | 18745 |

**Supplemental Table 3:** Gene ontology enrichment analysis for cellular content. Enrichment analysis for DEGs was run using https://david.ncifcrf.gov.

**Supplemental Movie 1:** A representative time lapse sequence shows profound contractile behavior in AOC cultures after introduction of fibroblasts. Large multicellular 3D nodules are aggregates of PANC1 cells formed over 7 days, and single cells are MRC5 normal human fibroblasts overlaid immediately prior to the start of video acquisition. Phase contrast images were acquired using a 10X objective with ten minutes between frames. Playback speed is 20 frames per second. This sequence moves to a different spatial position about 6 seconds into playback to show continued contraction.

**Supplemental Movie 2:** A representative time lapse sequence of PANC1 OR and MRC5 AOCs. The initial multicellular 3D nodules are aggregates of PANC1OR cells formed over 7 days, and single cells are MRC5 normal human fibroblasts overlaid immediately prior to the start of video acquisition. Phase contrast images were acquired using a 10X objective with ten minutes between frames. Playback speed is 20 frames per second.
